# Ancestral neural computations constrain the evolution of novel computations in simulated color vision networks

**DOI:** 10.1101/2019.12.30.891382

**Authors:** Nathan P. Buerkle, Stephanie E. Palmer

## Abstract

Efficient coding has been a successful organizational principle in neuroscience, but a more general theoretical framework needs to include the capacity for biological constraints to impede the realization of optimal circuit design. Here, we explore how evolution shapes the computational organization of a circuit using color vision as a model system. Taking a theoretical, machine learning approach allowed us to simulate the evolution of tetrachromatic color vision from a trichromatic ancestor both within and across distinct phylogenetic lineages. Analyzing network performance showed that trichromatic starting weights impose a significant constraint on learning rate, although the incremental increase in input layer complexity leads to better overall performance. Analyzing hidden layer computations showed that ancestry severely constrained evolution into a restricted and predictable portion of the theoretically available computational state space. Overall, our simulations of color vision evolution suggest that phylogenetic history is an important aspect of the functional organization of neural circuits.

## Introduction

Animals exhibit a rich diversity of behaviors and perceptual capacities that evolve as adaptive responses to ecological pressures. However, the evolutionary processes that give rise to behavioral variability do not necessarily occur as isolated, *de novo* searches for optimal solutions (Fawcett et al., 2013; Parker and Smith, 1990). Instead, factors such as phylogenetic history (Hale, 2014), development (Smith et al., 1985), and evolvability (Wagner and Altenberg, 1996) have the capacity to constrain and bias evolutionary trajectories. Accounting for these constraints and studying the mechanistic basis of adaptation has been invaluable for better understanding evolutionary patterns in morphological traits (Brakefield, 2011), protein sequences (Storz, 2016), and gene regulatory networks (Stern, 2013). Direct application of these concepts to the brain and behavior, in contrast, has been relatively limited (but see Ding et al., 2019; Katz, 2007; Seeholzer et al., 2018; Tosches, 2017).

Due to the complex and highly integrated organization of neural circuits, evolutionary constraints are thought to be especially strong for the brain and behavior (Katz, 2011; Tierney, 1995). Explicit and implicit assumptions of optimal neural coding are common throughout neuroscience, but ancestry and evolutionary constraints could potentially preclude this complex system from reaching a globally optimal solution. Instead, nervous systems may be biased into locally optimal solutions. A clear example comes from the well-studied jamming avoidance response in weakly electric fish, which is elegant in its computational implementation even though substantially simpler algorithms are theoretically possible (Heiligenberg and Rose, 1985; Rose, 2004). Comparative work further suggests the ancestral neural circuit organization was an important factor biasing evolution towards the observed, relatively complex computation (Matsubara and Heiligenberg, 1978; Rose and Canfield, 1991). Thus, an appreciation and consideration of phylogenetic history and its role in shaping neural circuit structure and function could lead to greater insights into principles defining how nervous systems process information and generate behavior.

Motivated by the diversity in photoreceptor spectral tuning across insects, color vision represents an attractive system for studying the evolution of neural computation. The ancestral insect eye most likely comprised ultraviolet (UV), blue, and green photoreceptors capable of trichromatic color vision (Briscoe and Chittka, 2001). A common adaptation, most notably in numerous butterflies, is the addition of a fourth, red-sensitive photoreceptor that expands color vision to tetrachromacy (Koshitaka et al., 2008; Stavenga and Arikawa, 2006; Zaccardi et al., 2006). Although evolving a new photoreceptor appears to be relatively simple (Briscoe, 2002; Frentiu et al., 2007), a novel photoreceptor alone may be insufficient to improve color vision and instead require additional changes to central circuits. No ecological or selective pressures have been identified to explain which insects do and do not have tetrachromatic vision, raising the possibility that evolutionary constraints influence and possibly impede color vision expansion (Briscoe and Chittka, 2001).

The relatively simple and well-described opponent coding mechanism underlying color vision makes it especially amenable to evolutionary questions. Opponent neurons compare different photoreceptor responses with spectrally antagonistic excitation and inhibition and arise early in visual circuits (Lee et al., 1989; Paulk et al., 2009). In theory, tri- and tetrachromatic vision require only 2 and 3 unique opponent channels, respectively (Chittka, 1992; Hurvich and Jameson, 1957; Vorobyev and Osorio, 1998), and this simplicity can facilitate comparisons across species (Fig. 1A). Moreover, despite being a low dimensional computation, the available computational state space is huge and theoretically infinite, as numerous unique combinations of opponent channel can lead to perceptually equivalent color discrimination (Chittka, 1992; Vorobyev and Osorio, 1998). This lack of constraint on how to implement opponent coding means that similarities and differences between independent origins of tetrachromatic vision can more confidently be ascribed to evolutionary rather than computational constraints.

**Figure 1:**
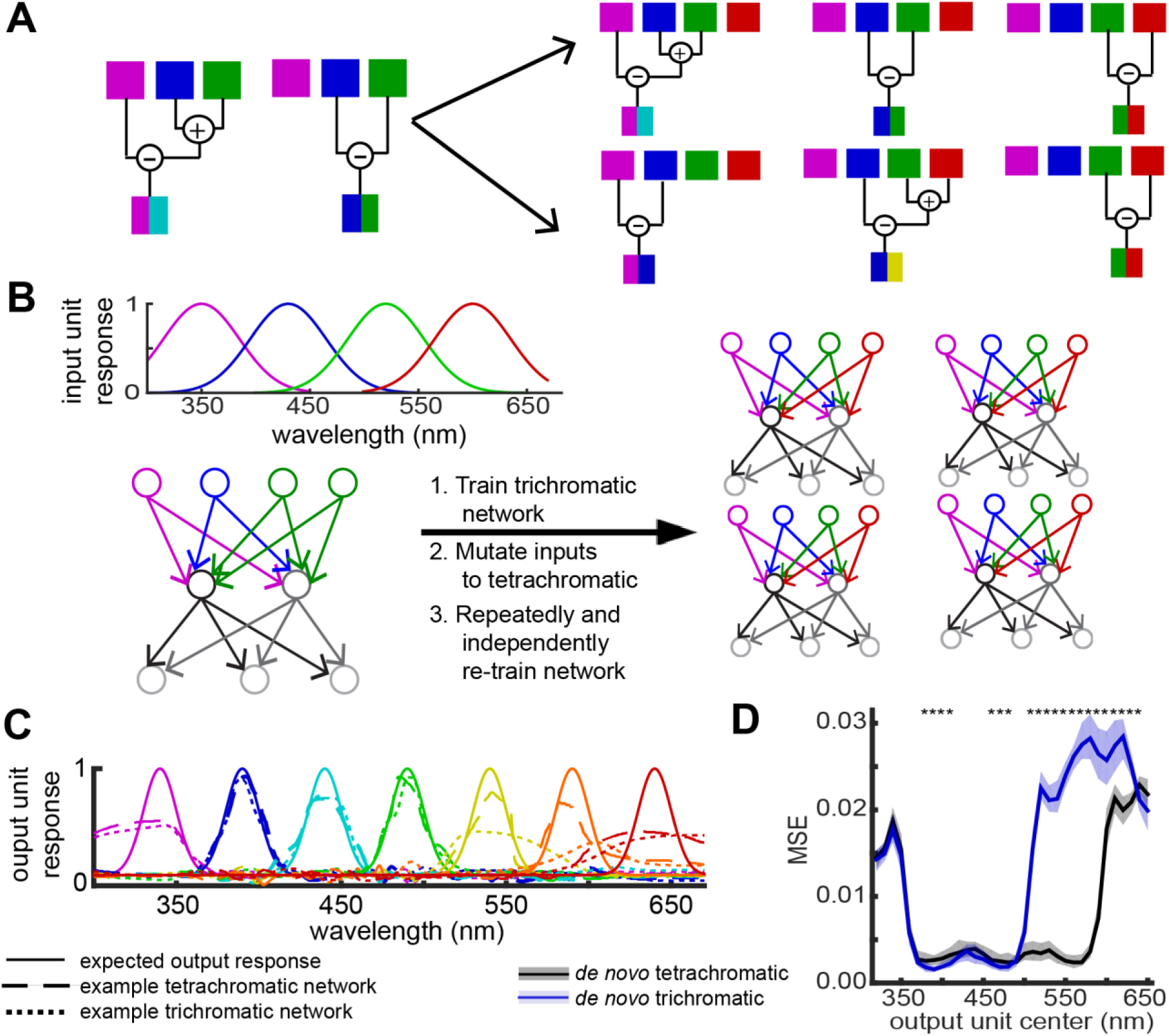
Network design. A. In theory, trichromatic color vision requires three photoreceptor types and two color opponent channels. The left shows one of numerous potential pairs of opponent channels, corresponding to UV vs. (B+G) and B vs. G. Tetrachromatic vision requires a fourth photoreceptor type and a third opponent channel (right). To evolve tetrachromatic computations from a trichromatic ancestor, a network could inherit the original channels and layer on a third (top) or evolve a completely novel set of opponent channels (bottom). B. Color vision evolution was simulated using feed-forward networks and a two-stage training procedure. Each network had 30 hidden units and 34 output units. Each of 100 trichromatic networks was re-trained 100 times, for 10,000 “evolved” networks. *De novo* networks trained for tetrachromacy directly from random starting weights were created for comparison. C. The output layer was a filter bank of narrowly tuned Gaussians that evenly tiled visual space. Shown here are the tuning curves for a representative trichromatic and tetrachromatic network. D. Network performance was measured as the MSE between the observed and expected tuning curve for each output unit. Performance as measured for tri- (n=100) and tetrachromatic (n=100) networks trained from random starting weights. Shading shows the 25^th^ and 75^th^ percentile across all networks. Asterisks indicate significant performance differences, with Cohen’s d > 0.5 used as an effect size based significance criterion.

For this study, we adopted a theoretical, machine learning approach to simulate the evolution of tetrachromatic vision from a trichromatic ancestor. By generating numerous trichromatic ancestors and numerous independent origins of tetrachromatic vision from each one, we were able to simulate convergent evolution both within and across distinct phylogenetic lineages.

Comparisons of network performance, the overarching computational structure of each network, and homologous hidden units across networks that either shared or did not share the same trichromatic ancestor allowed us to assess how phylogenetic history might constrain the evolution of neural computation. Overall, our simulation results were broadly consistent with existing behavioral and neurophysiology data, color vision theory, and conceptual ideas in evolutionary biology.

## Color Vision Model

Feed-forward, 3-layer neural networks were trained to discriminate monochromatic inputs using a standard backpropagation learning algorithm (Fig. 1, see methods for details). The input layer represented initial phototransduction in the eye with UV, blue, green, and red photoreceptors modeled as Gaussian filters on the input light spectrum (Fig. 1B). The output layer was a filter bank of narrowly tuned Gaussians wavelength “categories” (Fig. 1C) that mimicked wavelength selective neurons found in both insects (Kien and Menzel, 1977; Swihart, 1972) and primates (Schein and Desimone, 1990; Zeki, 1980). Networks had a single hidden layer with a sigmoid nonlinearity. Our analyses focused on networks with 30 hidden units, but results were qualitatively similar regardless of hidden layer size (Fig. S1). Networks responded to monochromatic light stimuli, with input layer responses scaled by a random, multiplicative luminance factor that removed stimulus brightness as a learnable cue (Fig. S1B).

Using this simple network design, we simulated color vision evolution using a two-stage training procedure (Fig. 1B). First, a network was initialized with random starting weights in all layers and pre-trained for trichromatic vision with an input layer that had UV, blue, and green photoreceptors. This trained network, mimicking a trichromatic ancestor, was then “mutated” by converting a subset of green photoreceptors to red, matching the known evolutionary history of butterfly photoreceptors (Briscoe, 2008). This mutated network was then retrained for tetrachromatic vision 100 unique times, which we viewed as biologically analogous to 100 species that independently evolved tetrachromatic vision from the same trichromatic ancestor. We generated and evolved 100 trichromatic networks to simulate distinct phylogenetic lineages (e.g. butterflies, spiders, birds, etc.), for a total of 10,000 evolved networks.

If ancestry constrains evolutionary trajectories, we expect evolved networks sharing the same trichromatic starting weights to be more similar than networks with different trichromatic starting weights. These potential constraints could be unique to a trichromatic ancestor, but they could also be more broadly applicable to any set of starting weights. To control for this possibility, we trained and analyzed an additional set of *de novo* networks. These networks lacked a pre-training step and were instead trained for tetrachromatic vision directly from random starting weights. Mirroring the evolved networks, we generated 100 sets of random starting weights and trained each starting point 100 unique times, again for a total of 10,000 *de novo* networks. Because of large sample sizes, nearly every statistical comparison was highly significant using a standard p value, so we instead adopted a stricter, effect-size based significance criterion of Cohen’s d > 0.5 for most analyses.

## Results

### Tetrachromatic networks had an expanded range of good color

We first validated our network design and training protocol by comparing the performance of *de novo* tri- and tetrachromatic networks trained from random starting weights. To evaluate performance, we constructed tuning curves for each output unit individually by measuring its response to monochromatic stimuli spanning the full visual spectrum and averaging across luminance factors (Fig. 1C). Performance was then defined as the mean squared error (MSE) between the expected and observed tuning curve. Tetrachromatic networks performed well across the entire visual spectrum, while trichromatic networks failed to learn for long wavelength stimuli (>500 nm, Fig. 1D) due to the lack of a red photoreceptor. Increasing hidden layer size did not improve long wavelength performance, but removing luminance variation eliminated differences between tri- and tetrachromatic networks (Fig. S1B).

Performance generally followed the Fisher Information of the input layer (Fig. S1A), which broadly matches behavioral discrimination thresholds measured in insects and vertebrates (Koshitaka et al., 2008; Neumeyer, 1986; Valois and Jacobs, 1968; von Helversen, 1972). That is, output units tuned to wavelengths where photoreceptor tuning curves intersect had the best performance, while performance decreased slightly between these peaks. Notably, trichromatic networks outperformed tetrachromatic networks at these Fisher Information peaks, likely reflecting a smaller range of learnable stimuli but an equal number of hidden units. Overall, the performance of these networks showed clear differences between tri- and tetrachromatic networks, which provided a clear set of comparison networks to ask how color vision may evolve.

### Mutating the input layer impaired network performance

To simulate the evolution of tetrachromatic vision, the input layer of trained trichromatic networks was “mutated” by converting a subset of green photoreceptors to red (Fig. 1B). This mutation, prior to any retraining, resulted in a mismatched tetrachromatic input layer and trichromatic hidden layer. This situation likely mirrors biology, where a novel photoreceptor necessarily precedes changes to central brain circuits. Output units tuned to short wavelengths were unaffected by the mutation, while performance was significantly impaired for middle and long wavelength output units (Fig. 2A). Interestingly, the mutation affected every network similarly (Fig. 2A, note error bar size), suggesting that changes to peripheral sensing caused similar disruptions to sensory perception regardless of the specific computations implemented by each network.

**Figure 2:**
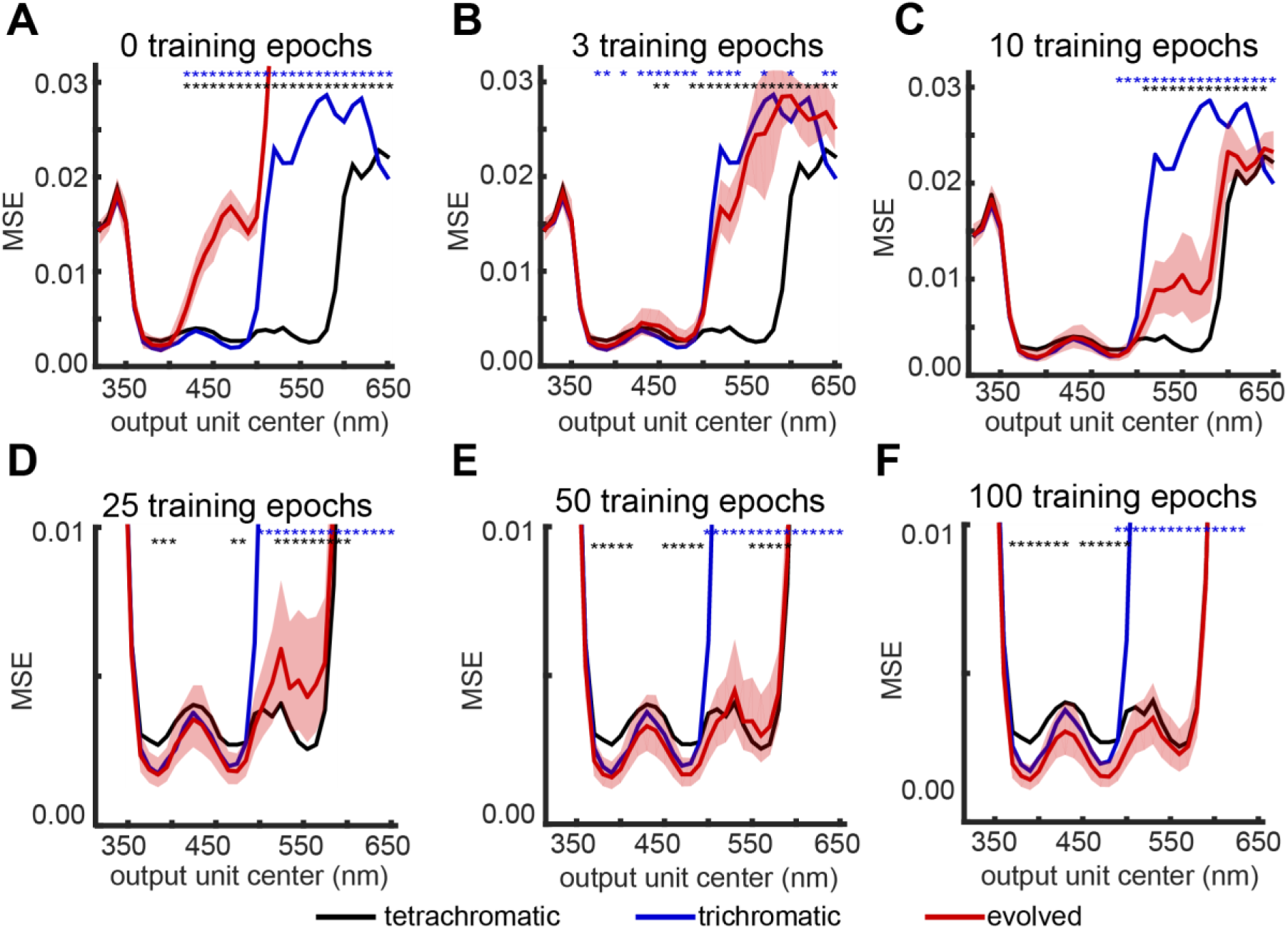
Evolved network performance over training time. The performance of evolved networks was tracked over training time and compared to *de novo* tri- and tetrachromatic networks trained from random starting weights after A) 0, B) 3, C) 10, D) 25, E) 50, and F) 100 training epochs. 50 training epochs matches the total training time of the *de novo* networks. Shading shows the 25^th^ and 75^th^ percentile and was omitted for *de novo* networks for clarity. Blue and black asterisks show significant differences between evolved and either tri- or tetrachromatic networks, respectively, with Cohen’s d > 0.5.

We further examined the cause of this impaired MSE performance by fitting Gaussians to each output tuning curve (Fig. S2). For tri- and tetrachromatic networks, tuning centers, widths, and amplitudes generally matched the target. In mutant networks, long wavelength output units became broadly tuned and were poorly fit by a Gaussian. However, middle wavelength output units typically maintained narrow tuning curves with a 5-10 nm shift in the center of tuning accounting for the majority of the impaired MSE (Fig. S2). These relatively minor shifts suggest the effect of mutation was smaller than the MSE metric initially showed. Instead, mutant networks likely retain trichromatic discrimination, although even small perceptual deficits could lead to selection against the expression of a novel photoreceptor.

### Evolved networks outperform *de novo* networks

Consistent with peripheral sensory systems being evolutionarily labile (Bendesky and Bargmann, 2011), mutant networks appeared to have only minor color vision deficits. However, a novel red photoreceptor alone was insufficient to expand color vision from tri- to tetrachromatic, indicating that further modifications to the circuit were necessary. Thus, we next asked if the networks were capable of adapting to a mutated input layer by retraining the mutant networks and tracking performance over training epochs (Fig. 2). These networks compensated for the new photoreceptor quickly, but improvement to tetrachromatic performance proceeded substantially slower. After 3-5 training epochs, the performance of these evolved networks broadly matched the original trichromatic network (Fig. 2B). After 50 training epochs, which matches the total training time of the *de novo* networks, evolved networks were effectively tetrachromatic, although long wavelength output unit performance remained slightly but significantly worse than *de novo* networks (Fig. 2E). Extending training to 100 epochs improved long wavelength performance to match *de novo* networks (Fig. 2F).

For short and middle wavelength output units, *de novo* trichromatic networks performed significantly better than *de novo* tetrachromatic networks (Fig. 2). Surprisingly, evolved networks inherited this improved performance, highlighting the potential value of curriculum learning (Bengio et al., 2009) in an evolutionary context. We investigated this effect further by examining the weights connecting the input photoreceptors to the hidden layer (Fig. S3). We defined the connection strength of a hidden unit as the sum of the absolute value of the four input weights (L1 norm). Random starting weights had consistently high connection strengths, and training led almost exclusively to a decrease. A tetrachromatic input layer led to a relatively uniform distribution of connection strengths. Interestingly, a trichromatic input layer led to significantly more reduction in connection strength. Evolved networks inherited this trichromatic distribution and reduced connection strength slightly further.

This extra decrease in connection strength for trichromatic networks was reminiscent of L2 regularization, which is commonly used in machine learning to prevent over-fitting. To ask how regularization affected network performance, we generated a new set of *de novo* tetrachromatic networks with an L2 regularization term explicitly added to the training protocol (Fig. S3). As expected, these networks had smaller connection strengths, but performance did not improve. Performance at Fisher Information peaks matched networks without regularization but was significantly worse between these peaks when regularization was included. Overall, these results show that incremental increases in network complexity can lead to improved performance.

### Different starting networks learn at different rates

Given a gradient descent learning algorithm, it was not surprising that every network successfully evolved tetrachromatic performance with enough training epochs. However, rather than strictly prohibiting an adaptive phenotype, evolutionary constraints are primarily thought to affect how accessible an adaptive phenotype is (Smith et al., 1985). For our networks, this could mean that different trichromatic networks may vary in how quickly they gain tetrachromatic performance. To investigate this possibility, we examined how different sets of starting weights influenced network performance after 10 training epochs (Figs. 2C, 3). For this and subsequent analyses, networks were separated into groups of 100 that either shared or did not share the same starting weights, resulting in four classes: 1) evolved networks with the same trichromatic starting weights, 2) evolved networks with different trichromatic starting weights, 3) *de novo* networks with the same random starting weights, and 4) *de novo* networks with different random starting weights. Each class comprised 100 unique groups with each group containing 100 networks.

**Figure 3:**
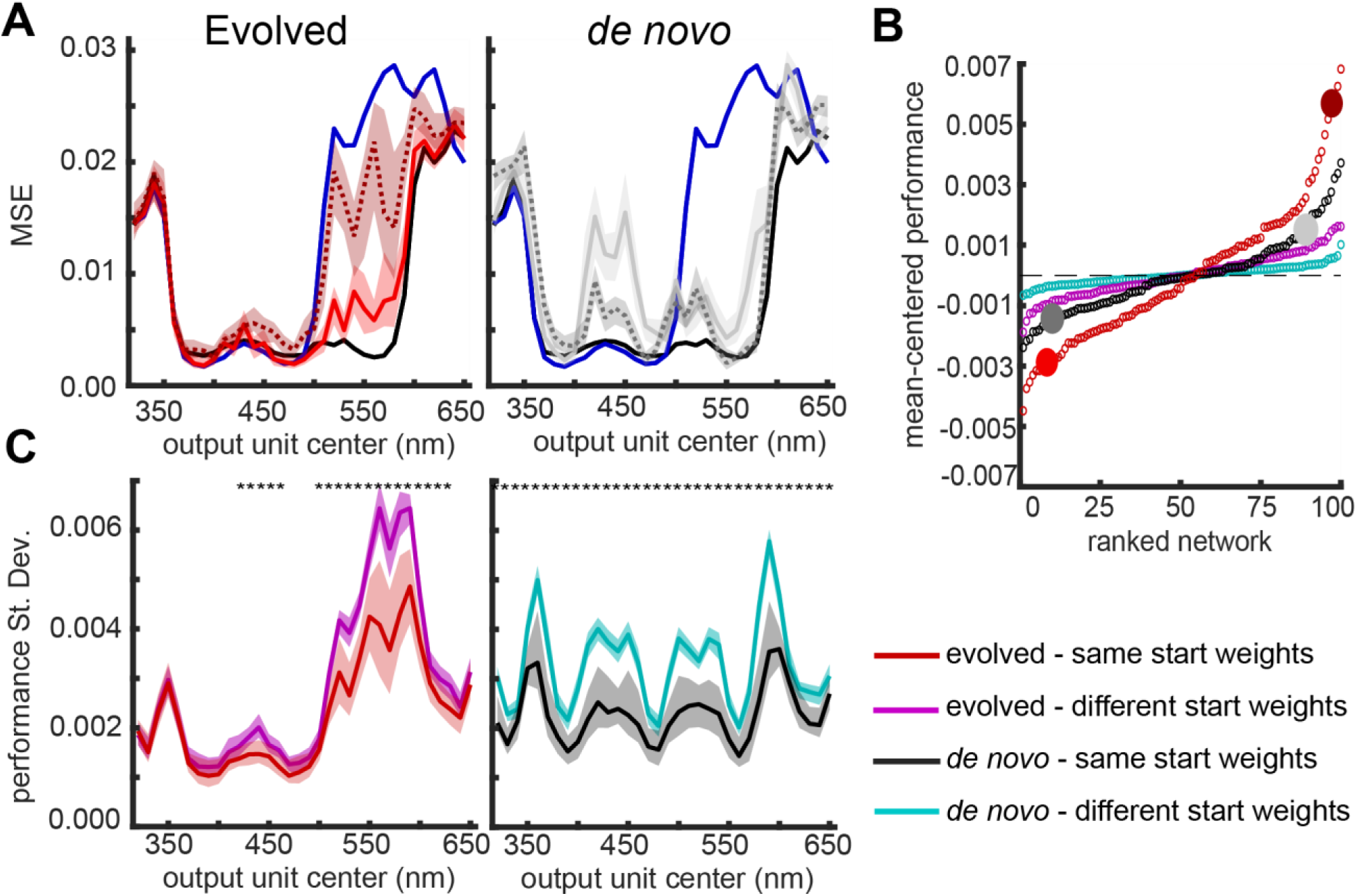
Starting weights influence learning rates. A. Networks were separated into groups of 100 that shared or did not share the same starting weights, and we analyzed performance after 10 training epochs for both evolved (left) and *de novo* (right) networks. Black and blue lines show the performance of fully trained tri- and tetrachromatic networks for reference. Solid lines show the performance of 100 networks that share the same starting weights. Dotted lines similarly show the performance of 100 networks that share a different set of the same starting weights. Shading represents the 25^th^ and 75^th^ percentile of performance across the 100 networks sharing the same starting weights. B. For each group of 100 networks that either shared or did not share the same starting weights, performance after 10 training epochs was assessed by averaging performance across all networks and long wavelength output units within a group. Distributions for each type of group were mean-centered and compared using a two-tailed Kolmogorov-Smirnov test. All pairwise comparisons were significantly different (p < 0.001). The large, filled circles correspond to the groups shown in panel A C. The standard deviation of performance was calculated for each output unit within a group of 100 networks. Shading shows the 25^th^ and 75^th^ percentile across the 100 groups. Asterisks denote significant differences with Cohen’s d > 0.5.

Visual inspection of network performance showed that starting weights influenced learning rate for both evolved and *de novo* networks. After 10 training epochs, evolved networks derived from some trichromatic starting weights approached tetrachromatic performance, while the descendants of other trichromatic networks showed minimal improvement beyond the original trichromatic network (Fig. 3A). For *de novo* networks, performance after 10 epochs at Fisher Information peaks tended to be good regardless of starting weights, while performance between these peaks were more dependent on the specific starting weights (Fig. 3A). This result was reminiscent of animal behavior, where discrimination thresholds near Fisher Information maxima are often similar across species, while thresholds between these regions of best discrimination vary more substantially (Koshitaka et al., 2008; Telles et al., 2016).

To quantify these visually observed effects, we compared the distribution of performance across the four network classes (Fig. 3B). For each group of 100 networks, performance after 10 training epochs was reduced to a single value by averaging across all networks and long wavelength output units (>500 nm). With 100 groups per class, this performance value was then compared across classes using a Kolmogorov-Smirnov test, which showed that every class was significantly different from the others (p < 0.001). Consistent with this effect, we also observed that within-group performance was less variable for groups that shared the same starting weights compared to groups of networks with different starting weights (Fig. 3C). Groups of evolved networks with the same trichromatic starting weights exhibited the largest variability in this performance metric (Fig. 3B), indicating that trichromatic starting weights had a strong influence on learning rate. From some trichromatic starting weights, tetrachromacy was easily accessible, while other starting weights impeded tetrachromacy evolution.

### Starting weights constrained the overarching computational structure of the hidden layer

Every network successfully evolved tetrachromacy, which we viewed as biologically analogous to convergent evolution both within and across distinct phylogenetic lineages. A convergent phenotype alone, however, is not sufficient to conclude that starting weights constrained the network. Instead, this might reflect similar selective pressures (i.e. the same training target and cost function). Learning rate differences hinted at an evolutionary constraint, but a rigorous test required assessing the degree of similarity in the underlying computational implementation of color vision across networks. Thus, we next turned to analyzing the hidden layer, focusing on the weights connecting the input layer to the hidden layer.

We first used principal components analysis (PCA) to analyze the overarching structure of the hidden layer. Consistent with color vision theory (Chittka, 1992; Hurvich and Jameson, 1957; Vorobyev and Osorio, 1998), PCA reduced the dimensionality of the hidden layer from 30 hidden units to 3 color opponent channels (i.e. eigenvectors) that described 48.5 ± 12.2, 27.6 ± 5.5, and 16.0 ± 5.7 percent of the variance (92.1% in total) in hidden unit computations. Each channel had a UV, blue, green, and red component that consistently showed opponent interactions, indicated by a combination of positive and negative input weights (Fig. 4A). To assess computational similarities between networks, we concatenated the three opponent channels into a single 12-dimensional vector and performed hierarchical clustering on groups of networks that either shared or did not share the same starting weights (Fig. 4).

**Figure 4:**
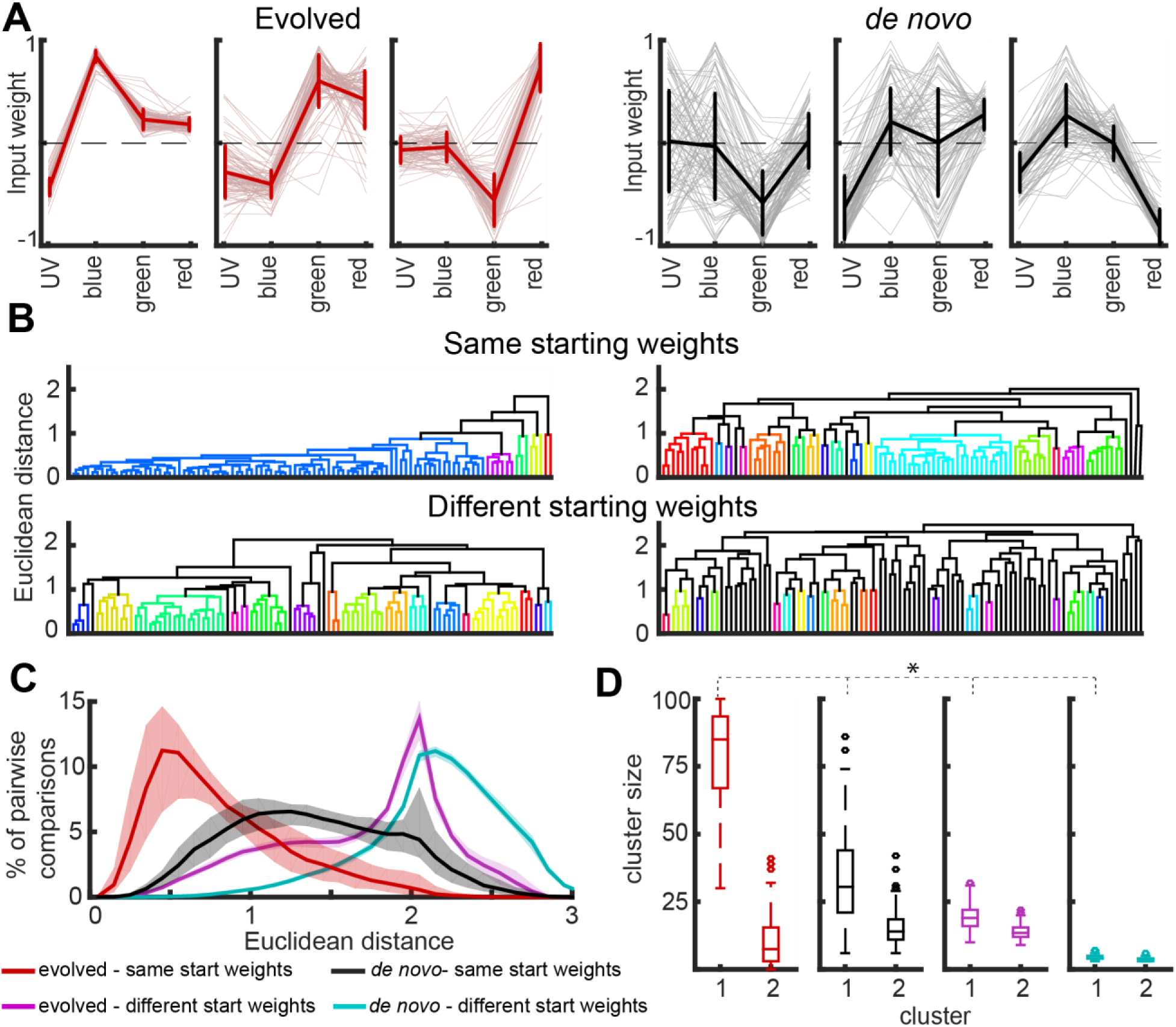
Starting weights constrain the computational structure of the hidden layer. A. PCA on the weights connecting the input layer to the hidden layer revealed 3 opponent channels (i.e. eigenvectors) that described the overarching computational structure of a network. Input weights with opposite signs are indicative of an antagonistic opponent interaction between photoreceptor types. The left three panels show the opponent channels for 100 evolved networks derived from the same trichromatic starting point. The right three panels similarly show the opponent channels for 100 *de novo* networks derived from the same random staring weights. Thick lines and error bars are mean **±** standard deviation. B. Opponent channels for groups of 100 networks that either shared or did not share the same starting weights were analyzed using hierarchical clustering. Depicted are representative dendrograms for each of the four classes of networks. The top row corresponds to the networks shown in panel A. Different colors represent different clusters with a clustering threshold of 1.0. C. Euclidean distances between opponent channels were measured for every pair of networks within a group of 100 and binned in intervals of 0.1. Shading shows the 25^th^ and 75^th^ percentile. D. Groups of networks were clustered using a distance threshold of 1.0. Boxplots show the distribution of cluster sizes for the two largest clusters for each class of groups.

Comparing groups of *de novo* networks that shared the same random starting weights with groups that did not showed that starting weights constrain and bias learning trajectories. *De novo* networks with different starting weights used a diverse set of opponent channels, as the largest cluster contained between 3 and 7 out of 100 networks within a group (Fig. 4D). Moreover, the distribution of all pairwise Euclidean distances within a group was large and closely matched a null distribution generated from random numbers (Fig. 4C, S4, Jensen-Shannon Divergence = 0.02). The opponent channels for networks sharing the same random starting weights, in contrast, had a pairwise distance distribution shifted substantially towards 0 (Fig. 4C, JSD = 0.31) and an average of 33.8 ± 17.1 networks in the largest cluster (Fig. 4D). These results showed that although starting weights do constrain learning, *de novo* networks do retain a degree of diversity in how they implement tetrachromatic computations.

Trichromatic starting weights constrained computations significantly more than random starting weights. Groups of evolved networks sharing the same starting weights typically converged on just one or two computational motifs. Pairwise distances were shifted further towards 0 compared to *de novo* networks with the same starting weights (Fig. 4C, JSD = 0.19), and the largest cluster contained 79.2 ± 17.7 networks (Fig. 4D, Cohen’s d = 1.9). Only 2.5 ± 2.1 clusters were needed to account for 90 out of the 100 evolved networks in a group, whereas *de novo* networks required 12.3 ± 7.9 clusters (Cohen’s d = 1.7). Together, these results showed that starting weights constrained network computations, and trichromatic weights imposed especially strong constraints that biased networks into a severely restricted region of the available computational state space.

### Evolved network opponent channels are predictable

Groups of evolved networks with different trichromatic starting weights were broadly similar to groups of *de novo* networks with the same starting weights. The largest cluster had 19.4 ± 4.6 networks, and 13.7 ± 2.9 clusters were required to account for 90 networks. Interestingly, the pairwise distance distribution matched the distance distribution of the original trichromatic networks, which is essentially a group of 100 networks with different starting weights (Fig. S4B, JSD = 0.01). This similarity was surprising because one might assume that the inherently smaller dimensionality of trichromatic networks, which have only 2 opponent channels with three input weights, would lead to smaller distances. To understand how this similarity arose, we analyzed each opponent channel individually rather than as the concatenated group of three (Fig S4D).

We first noticed that the new, third channel for nearly every evolved network had green vs. red opponency regardless of the specific trichromatic starting weights (Fig. S4A). Hierarchical clustering of the third channel confirmed this observation (Fig. S4D). In contrast, the first and second channel matched previous results (Fig. 4), with groups sharing the same starting weights more similar than groups with different starting weights. We then calculated the Euclidean distance between these two opponent channels and their original trichromatic channels. These distances (0.33 ± 0.20) were significantly different from 0 suggesting that the channels were not static, but were also significantly smaller than *de novo* channel distances (0.55 ± 0.20, Cohen’s d = 1.1) or evolved networks with shuffled starting channels (0.66 ± 0.28, Cohen’s d = 1.3).

Overall, these results indicated that networks evolve tetrachromatic vision by largely inheriting the original two opponent channels and adding a third, orthogonal channel specifically implementing green vs. red color opponency. This result is consistent with how networks performed, as limited disruption to short wavelength computations led to limited disruptions in short wavelength performance. Thus, maintenance of this performance may play a causal role in biasing tetrachromatic computations. *De novo* networks, in contrast, were more free to vary and find unique computational solutions that spanned the full computational state space.

### Evolved and *de novo* hidden units learn in different ways

The opponent channels revealed by PCA describe the overarching algorithms used for color discrimination in our networks. These algorithms, in turn, were implemented by a population of individual hidden units that were the direct target of training. To understand how the tuning of these hidden units changed to support color computation, we compared the tuning of each hidden unit before and after training using two metrics (Fig. 5A). The cosine distance measured changes in opponent tuning, and the city block distance measured changes in connection strength. Moreover, because each starting network was independently trained 100 times, each hidden unit had a distribution of distances (Fig. 5B). Comparing the tuning of these homologous hidden units then allowed us to determine whether similar opponent channels emerge due to similar changes in individual hidden units.

**Figure 5:**
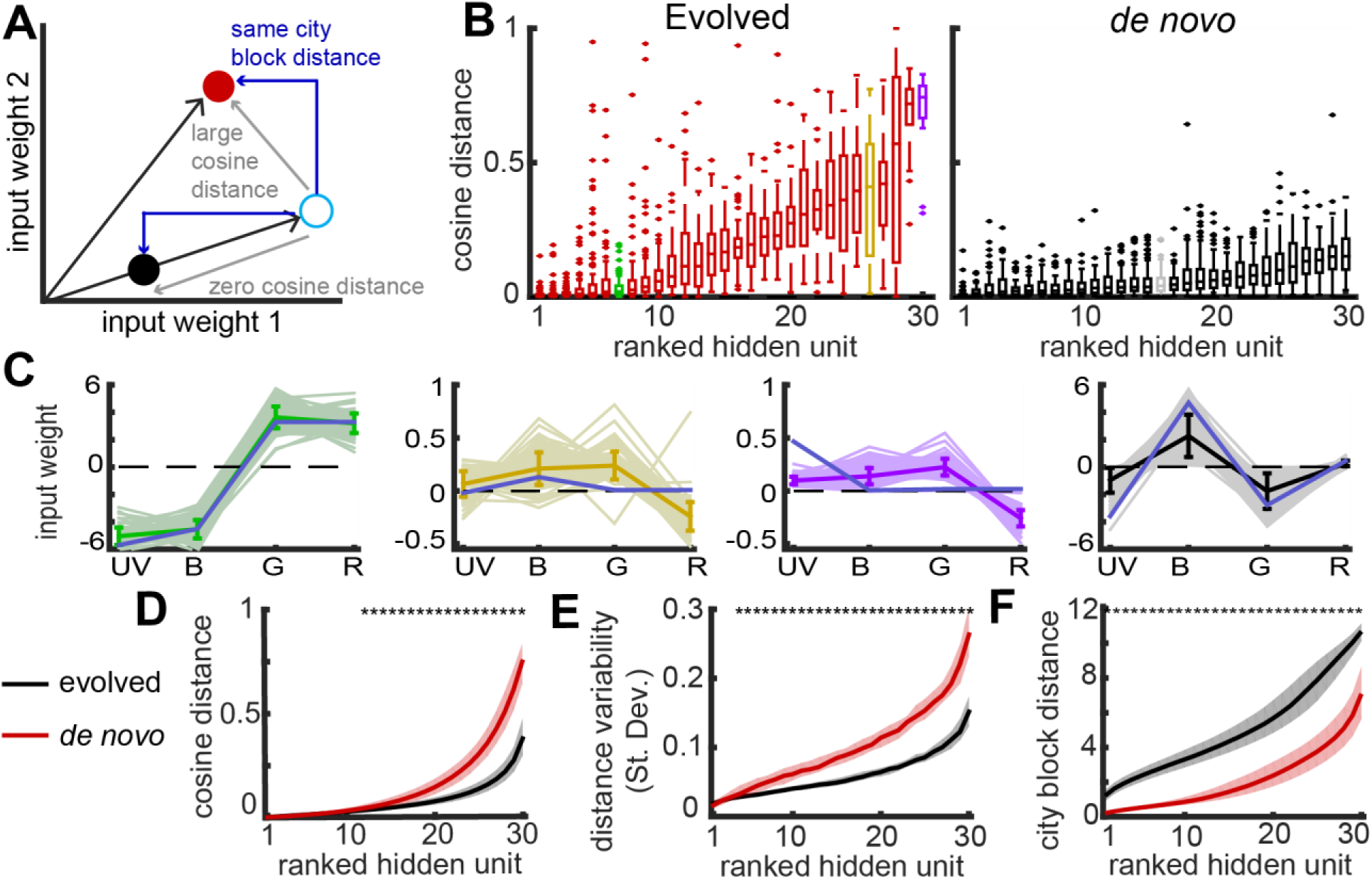
Training leads to larger changes in the tuning of evolved network hidden units. A. Schematic diagram shows the two types of distance used to measure hidden unit changes. The total length of the blue arrows is the city block distance, and the distance of each filled circle to the open circle is the same. The cosine distance measures the relative change in input weights while ignoring differences in overall weight magnitude. Because the black circle and open circle lie on the same vector, the cosine distance is 0, while the distance between red and the open circle is large. B. Tuning changes for each hidden unit were measured as the cosine distance between the input weights of a hidden unit before training and the input weights after training. Boxplots show the distribution of cosine distances for 100 evolved (left) and *de novo* (right) networks derived from the same starting weights and ranked in order of median distance. Colored boxes correspond to plots in panel C. C. Each panel shows the tuning of an example hidden unit, with colors corresponding to the colored boxes in panel B. The solid line and error bars are the mean ± standard deviation. The blue line shows the tuning of the original, untrained hidden unit. Note differences in the scale of the y-axis. D. Cosine distance distributions were measured for each of the 100 starting networks and ranked in order of increasing cosine distance. Shading shows the 25^th^ and 75^th^ percentile. Asterisks show significant differences between evolved and *de novo* networks with Cohen’s d > 0.5. E. Same as D, but rather than median cosine distance, the standard deviation of the cosine distance distribution was measured. F. Same as E, but instead of measuring the cosine distance, here we measured the city block distance.

Surprisingly, in contrast to opponent channel results, evolved network cosine distances were significantly larger (Fig. 5D) and more variable (Fig. 5E) than *de novo* networks. Approximately 10 hidden units per network had especially large cosine distances. These 10 hidden units typically had small connection strengths and weak opponency before training and a clear red vs. green opponent interaction after training (Fig. 5C, yellow and purple). Hidden units with the smallest cosine distances usually had large connection strengths and strong opponent interactions (Fig. 5C, green). Interestingly, evolved networks with 50 hidden units similarly had approximately 10 hidden units with large distances (Fig. S5). These results suggest the tuning of most hidden units were rigidly constrained to support ongoing short and middle wavelength color vision, while a select group was flexible and supported improved color vision through the development of green vs. red opponency.

Rather than changing opponent tuning, *de novo* networks instead learned primarily by modifying connection strength through the proportional scaling of all input weights. Cosine distances were small, showing that hidden unit opponent tuning changed little (Fig. 5E). City block distances, in contrast, were significantly larger for *de novo* networks, showing that they changed their connection strengths (Fig. 5F). Further, the connection strength of homologous hidden units varied more for *de novo* networks compared to evolved networks (Fig. S6). These results are broadly consistent with color vision theory (Chittka, 1992; Vorobyev and Osorio, 1998), suggesting that any randomly generated set of opponent cells is capable of good discrimination, with connection strength changes potentially useful for matching network specific details such as output unit amplitude. Together with the cosine distance analysis, these results indicate that evolved and *de novo* networks differ in which features of the connection pattern are flexible and change to support learning.

Given these differences in learning, we next wanted to ask about the relative contribution of opponent tuning and connection strength to opponent channel similarities. To investigate this, we rescaled the connection strength of each hidden unit to a unit vector to remove connection strength variability and re-performed the opponent channel, PCA analysis (Fig. S6). This normalization increased opponent channel similarities for both evolved and *de novo* networks, but evolved networks remained significantly more similar. Thus, despite *de novo* network cosine distances being small, the minor opponent tuning changes combined across the full population of 30 hidden units contributed significantly to the observed differences in the overarching computational structure of the hidden layer.

### Hidden unit outputs are robust to input variability

In addition to large cosine distances, evolved network hidden units also exhibited a high degree of cosine distance variability (Fig. 5E). These differences in the opponent tuning of homologous hidden units could make important contributions to how different networks function, but they could also reflect network computations that are robust to input variation. Prior analyses focused on the weights connecting the input photoreceptors to the hidden layer, so to distinguish between these possibilities, we now turned to examining the output response of a hidden unit. Importantly, the response of a hidden unit is passed through a sigmoid nonlinearity before reaching the output layer, which could potentially make its response relatively insensitive to precise input weights while maintaining sensitivity to the sign of the input.

To measure the output response curve of each hidden unit, we calculated its response to monochromatic stimuli (Fig. 6A) and transformed that response using a sigmoid nonlinearity (Fig. 6B). Hidden units typically responded to a stimulus with a binary 0 or 1 response, with relatively sharp transitions between ‘on’ and ‘off’. In general, response curves for homologous hidden units were variable prior to the nonlinearity and became more similar after the nonlinearity. The exception to this effect, however, occurred when an input weight to a pair of homologous hidden units had different signs. In these instances, even small differences in input weight could lead to large differences in output response due to the nonlinearity (Fig 6).

**Figure 6:**
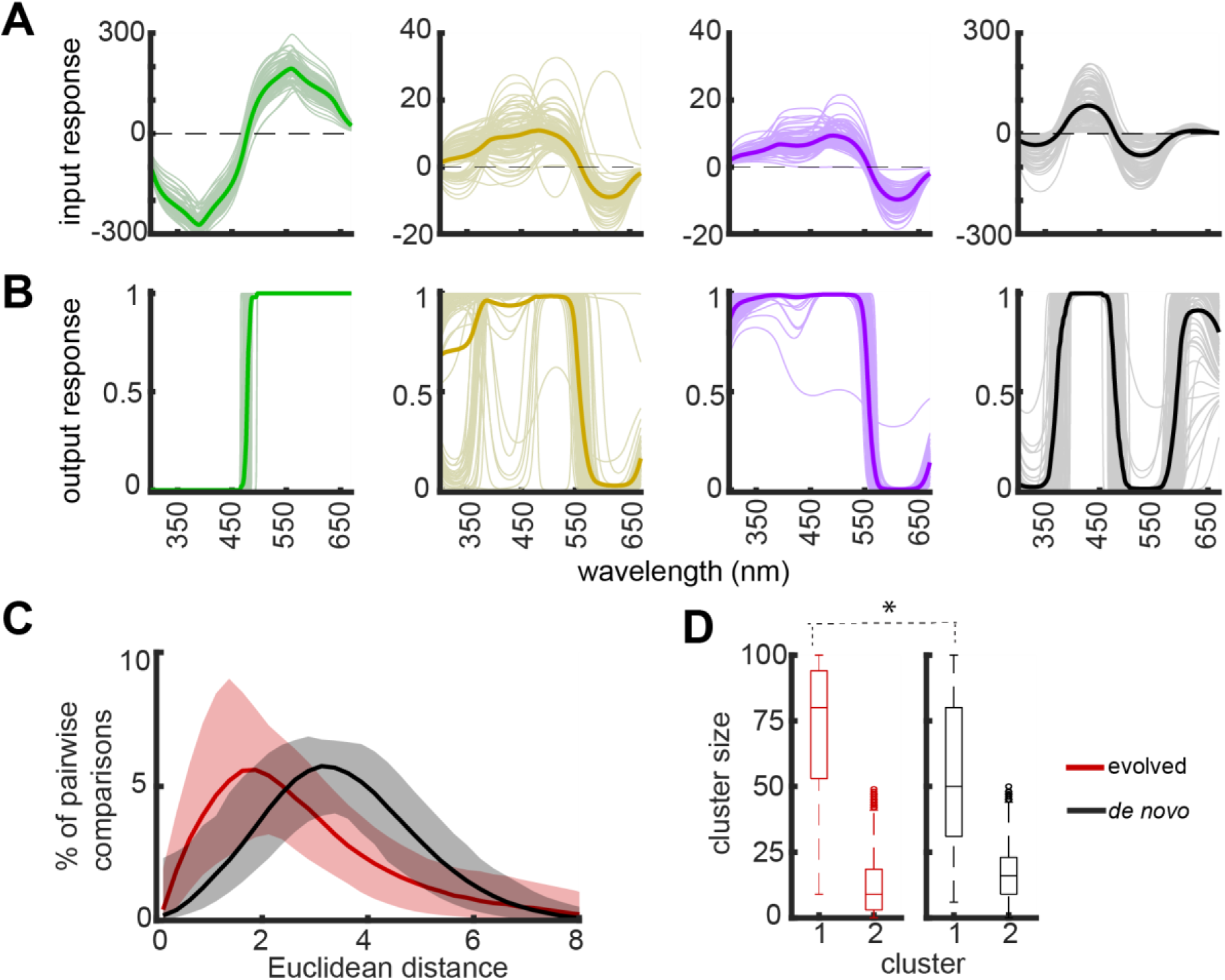
Hidden unit nonlinearity limits the effect of opponent tuning changes. A. The input response of a hidden unit to a monochromatic stimulus was measured as the activation of the input photoreceptors convolved with its input weights. Colors correspond to the hidden units shown in Figure 5. B. The output response curve of a hidden unit is calculated as the input response passed through a sigmoid nonlinearity, which limits the response to values between 0 and 1. C. The output response curves of homologous hidden units were analyzed using hierarchical clustering. The distribution of pairwise distances shows a shift towards 0 for evolved networks across all hidden units (100 starting networks X 30 hidden units per network. Shading shows the 25^th^ and 75^th^ percentile. D. Output response curves were clustered with a distance threshold of 3.0. The largest cluster for evolved network hidden units was significantly larger than *de novo* networks (Cohen’s d = 0.7)

We quantified these observations using hierarchical clustering and compared evolved to *de novo* networks (Fig. 6). Across all sets of homologous hidden units (100 starting networks X 30 hidden units per network), output response curves were more similar for evolved network hidden units. The distribution of pairwise distances was shifted slightly towards 0 for evolved networks (Fig. 6C, JSD = 0.05), which led to a significant increase in the size of the largest cluster (Fig. 6D). Overall, this result shows that the previously observed diversity in opponent tuning has a relatively minor effect on network function. This capacity to modify weights without disrupting network function may facilitate evolved network learning by allowing the network to flexibly enter and explore new computational states with minimal effects on the existing computational structure of the network.

## Discussion

Phylogenetic history has the capacity to constrain and bias future evolutionary trajectories, and these effects may be especially strong for complex neural circuits and the computations they implement (Katz, 2011; Smith et al., 1985). To examine the role of ancestry on neural computation, we simulated the convergent evolution of tetrachromatic vision both within and across distinct phylogenetic lineages using a simple neural network and training protocol. In our networks, trichromatic ancestry affected the rate of learning (Fig. 3) and also biased color vision computations into a restricted range of the available computational state space (Fig. 4). Overall, our results broadly show that trichromatic ancestry constrains the evolution of tetrachromatic vision, which can potentially explain why there is no clear ecological explanation describing which butterflies do and do not have tetrachromatic vision (Briscoe and Chittka, 2001). These findings suggest that evolutionary history is an important factor that influences the functional organization of neural circuits.

Expanding color vision from trichromatic to tetrachromatic necessarily begins with the evolution of a new photoreceptor. Consistent with the sensory periphery being evolutionarily labile and relatively immune to deleterious pleiotropic effects (Bendesky and Bargmann, 2011; Tosches, 2017), mutating the input layer led to minimal performance impairments in our model (Fig. 2,S2). Output unit tuning shifted, but the output layer maintained a structured organization of narrowly tuned units likely capable of trichromatic vision. Additionally, improved luminance discrimination can potentially offset any color vision impairments (Mancuso et al., 2009). Thus, a peripheral adaptation that creates a mismatched tetrachromatic eye and trichromatic brain is unlikely to play an important role in facilitating or impeding evolution.

Peripheral adaptation did not impair color vision but it also did not improve it, effectively making the mutation a neutral variant with respect to fitness. Neutral variants present at low frequency in a population are more likely to go extinct than drift to fixation, so the rate of circuit modification is likely an important factor determining whether a population retains a red photoreceptor. Every network successfully evolved tetrachromacy, but the rate of adaptation varied significantly between different trichromatic networks (Fig. 3). Fast learning networks convert the red photoreceptor from a neutral to advantageous variant quickly, possibly increasing the chance that it would survive in a real biological and ecological system.

Due to the nonlinear transformation of the hidden layer, hidden units input weights could change substantially with limited effect to the output response, and this flexibility may have contributed to diverse learning rates. Sign changes on the input weights were the major exception and was a necessary component for evolved networks to develop green vs. red opponency. This computational dynamic could have facilitated learning by allowing the network to make precise and targeted changes to long wavelength computations with minimal deleterious, pleiotropic effects on existing short and middle wavelength computations. Although we were unable to explicitly test this idea directly, hidden units in different trichromatic networks may vary in how flexible they are. If a network was less capable of decoupling the targeted development of green vs. red opponency from existing trichromatic computations, tetrachromacy evolution could become difficult and slow.

Despite the computational flexibility of single hidden units, starting weights strongly constrained the overarching opponent channel computations of a network. Consistent with color vision theory (Chittka, 1992; Hurvich and Jameson, 1957; Vorobyev and Osorio, 1998), *de novo* networks used diverse opponent channels that spanned an effectively infinite computational state space. Evolved networks, in contrast, were generally restricted to the original trichromatic channels and the addition of a specific green vs. red channel. Comparing networks with different trichromatic ancestors would then give the appearance of unconstrained computational flexibility. However, accounting for ancestry showed that the evolution was highly constrained. This is similar to hemoglobin evolution in high altitude birds, where protein sequence variability across independent evolutionary events suggests many paths to adaptation (Natarajan et al., 2016). However, accounting for phylogenetic history and existing neutral sequence variability revealed a constrained pattern of evolution.

Networks learned using a series of discrete training epochs comprised of numerous monochromatic stimuli. How individual training epochs relate to developmental or generational time in real species could vary substantially between different taxa. In mammals, activity dependent development plays a significant role in the functional organization of the cortex, so the entirety of training time could occur within single individuals, which is supported by experimental data in two species. Squirrel monkeys have two opsin genes, with the X-linked opsin having a green and red allele. Males and homozygous females are obligate dichromats, but because of X-inactivation, heterozygous females have both green and red photoreceptors. Behavioral tests have indirectly shown evidence of trichromatic color vision in these females (Jacobs, 2009). Rodents do not have natural allelic variation for the X-linked opsin, but experimental introduction of a red allele led to similar behavioral results (Jacobs et al., 2007), although the computational basis of this behavior and whether these animals exhibited true trichromatic vision remains uncertain (Makous, 2007). Nonetheless, genetics do have a significant role in behavior, and its interaction with activity dependent development remains unclear.

In contrast to the mammalian cortex, development of the insect nervous system appears to be more strongly genetically hardwired (Brochtrup and Hummel, 2011; Hiesinger et al., 2006). For these animals, each training epoch might correspond to a single adaptive mutation occurring over the course of generations. Single training epochs had variable effects on performance, ranging continuously from small to large, which mirrors the exponential distribution of effect size that genetic loci have on behavioral traits. While genetically specified wiring is dominant in insects, a degree of activity dependent development has been observed in *Drosophila* (Golovin et al., 2019; Kaneko et al., 2017; Vonhoff and Keshishian, 2017). The extent and function remain unclear, but the effect does appear to primarily affect connection strength rather than circuit structure. This limited plasticity could relate to the first training epochs, where trichromatic performance is restored in the absence of substantial changes to circuit computation.

In theory, color vision has an infinite number of perceptually equivalent computational solutions built through combinations of distinct and independent opponent channels (Chittka, 1992; Vorobyev and Osorio, 1998). This combination of a simple computation and a large state space made color vision a good model for evolutionary questions as it allowed for clear comparisons across networks. The lack of a globally optimal solution further meant that all of our results showing similarities between networks sharing the same starting weights can be attributed to network ancestry rather than constraints on the specific implementation of color vision. One limitation, however, is that this relatively simple state space is unlikely to be representative of all neural circuits and behaviors. Instead, other circuits likely have complex computational state spaces with numerous local minima that vary in performance and are separated by barriers that vary in how difficult they are to traverse. These rugged landscapes could introduce a complex and dynamic interaction between optimality and constraint. Having shown here that phylogenetic history does influence network evolution and computation, identifying a more complex computation suitable for evolutionary modeling would be valuable for investigating this relationship.

## Acknowledgements

We would like to thank Joe Lombardo, Ben Hoshal, and Siwei Wang for useful comments and discussion on previous versions of this manuscript. This work was supported by the University of Chicago Big Ideas Generator, the Alfred P. Sloan Foundation (SEP), and an NSF CAREER award 1652617 (SEP).

## Author Contributions

NPB conceived of the project, generated the models, and completed network analysis and comparisons. SEP supervised project development and analysis. NPB and SEP wrote the paper together.

## Methods

### Network design

Color vision networks were designed as 3 layer, feed forward networks that responded to monochromatic light stimuli. Preliminary networks were generated using a broad array of tuning parameters for both the input and output layers. The particular set of tuning parameters used in this study was chosen because it maximized performance differences between tri- and tetrachromatic networks. The input layer, simulating photoreceptors, had Gaussian tuning curves with 50 nm standard deviations centered on 350 (UV), 430 (blue), 520 (green), and 600 nm (red). Responses were scaled by a luminance factor between 1 and 100 and passed through a saturating nonlinearity that capped the response at 50. A total of 34 output units evenly tiled the visual range, from 320 to 650 nm in 10 nm increments. Each output unit had a standard deviation of 7.5 nm. The hidden layer had between 10 and 50 hidden units, with analyses focused on 30 hidden units. Hidden unit responses were passed through a sigmoid nonlinearity.

Networks were trained with the Levenberg-Marquardt backpropagation algorithm (Hagan and Menhaj, 1994), which is the Matlab default. Alternative learning algorithms failed to perform well. Training samples were monochromatic light stimuli paired with a luminance factor.

Wavelengths varied from 300 to 670 nm in 5 nm increments, and luminance factors varied continuously from 1 to 100. Each training epoch had 400 novel pairs of wavelength and luminance factor. Using this large training set minimized the influence of training data on learning trajectories and effectively isolated the effect of starting weights. Severely reduced training sets with 10 nm wavelength increments and 6 discrete luminance factors (228 total stimuli) resulted in only minor performance decrements.

Circuit evolution was simulated using a two-stage training process. First, we trained trichromatic networks with a UV, blue, and two green input unit units. Starting weights were set with the Nguyen-Widrow initialization algorithm (Nguyen and Widrow, 1990). The two green input units had identical starting weights, making them function as a single input. After 50 training epochs, one of the green input units was mutated to red, and the network was trained for 100 more epochs. The un-mutated, trichromatic network was also trained for 50 more epochs (100 total), and these networks were used for all analyses. We trained 100 trichromatic networks using different random starting weights. Each trichromatic network was independently evolved 100 times using different sets of training data, for a total of 10,000 networks. In preliminary analyses we compared networks with different starting weights and the same training data. Results did not differ from networks with different starting weights and different training data.

As a comparison for evolved networks, we also trained a set of *de novo* networks. These networks were not pre-trained with a trichromatic input layer, but were otherwise identical to the evolved networks. Instead, these networks were directly trained for tetrachromatic vision from random starting weights. Mirroring the evolved networks, we generated 100 sets of random starting weights, again using the Nguyen-Widrow initialization algorithm. Each random network was trained 100 independent times using a tetrachromatic input layer. All analyses on these networks were performed in the same way as the evolved networks.

### Network performance

We measured tuning curves for each of the 34 output units and compared them to the target response. Output unit responses were calculated for stimuli between 300 and 670 nm in 1 nm increments. Tuning curves were made by averaging the response across 10 luminance factors ranging from 10 to 100 in steps of 10. Performance was then defined as the MSE between the observed and expected tuning curve. Results were qualitatively similar when calculating the MSE for any single luminance factor. Overall performance decreased slightly without averaging, but the shape of the output units tuning curves was unchanged.

### Opponent channels

The overarching computational structure of the hidden layer was analyzed by using PCA to extract the opponent channels. For this analysis, each hidden unit was an observation, and the UV, blue, green, and red input weights were the variables. PCA reduced the dimensionality to 4 eigenvectors, the first three of which consistently had color opponent interactions. The fourth typically had all positive input weights indicating a luminance channel and was not analyzed further. All opponent channels were scaled to unit vectors to make them comparable, but the sign was arbitrary. To account for this, we calculated every pairwise correlation between an opponent channel for one network and all other networks in a group of 100. Opponent channels with at least half of the correlations less than zero had its sign flipped. We tried several different sign flipping procedures, and this correlation-based method maximized similarities between networks.

Similarities and differences between networks were measured using agglomerative hierarchical clustering. We compared both single channels and the full set of three opponent channels concatenated together. We also generated groups of random opponent channels as a null hypothesis. These opponent channels were made with a random number generator, scaled to a unit vector, and sign flipped using the same correlation procedure. There was no prior expectation on how the opponent channels should be ordered, and efforts to maximize similarities found no procedure better than ordering and labeling opponent channels according to percent of variance explained. We clustered networks using Euclidean distances between networks and a complete linkage algorithm, which maximized cophenetic correlations (c > 0.85). The clustering threshold was set at 1.0, but results did not depend on the precise threshold. In addition to cluster size, we also examined the distribution of all pairwise distances within a group of 100 networks. Pairwise distances were binned in distance intervals of 0.1 for visualization purposes and intervals of 0.01 for analysis since this bin size maximized distribution entropy.

The distributions of pairwise distances were compared between group types using the Jensen-Shannon divergence, a metric related to the Kullback-Leibler divergence (Amari et al., 1987).

### Hidden unit analysis

Homologous hidden units were compared between networks that shared the same set of starting weights. For cosine and city block distances, the input weights from each of the input layer photoreceptors were used to create a 4-dimensional representation of the hidden unit tuning. To generate tuning curves, hidden units responses were measured for monochromatic stimuli ranging from 300 to 670 nm in 1 nm increments and averaged over 10 luminance factors. For the input tuning curve, the input layer response to a stimulus was convolved with the input weights to a hidden unit. To generate the output tuning curve, these input tuning curves were passed through a sigmoid nonlinearity. Output tuning curves were compared and clustered between homologous hidden units using Euclidean distances and a complete linkage function.

---

**Figure S1.**
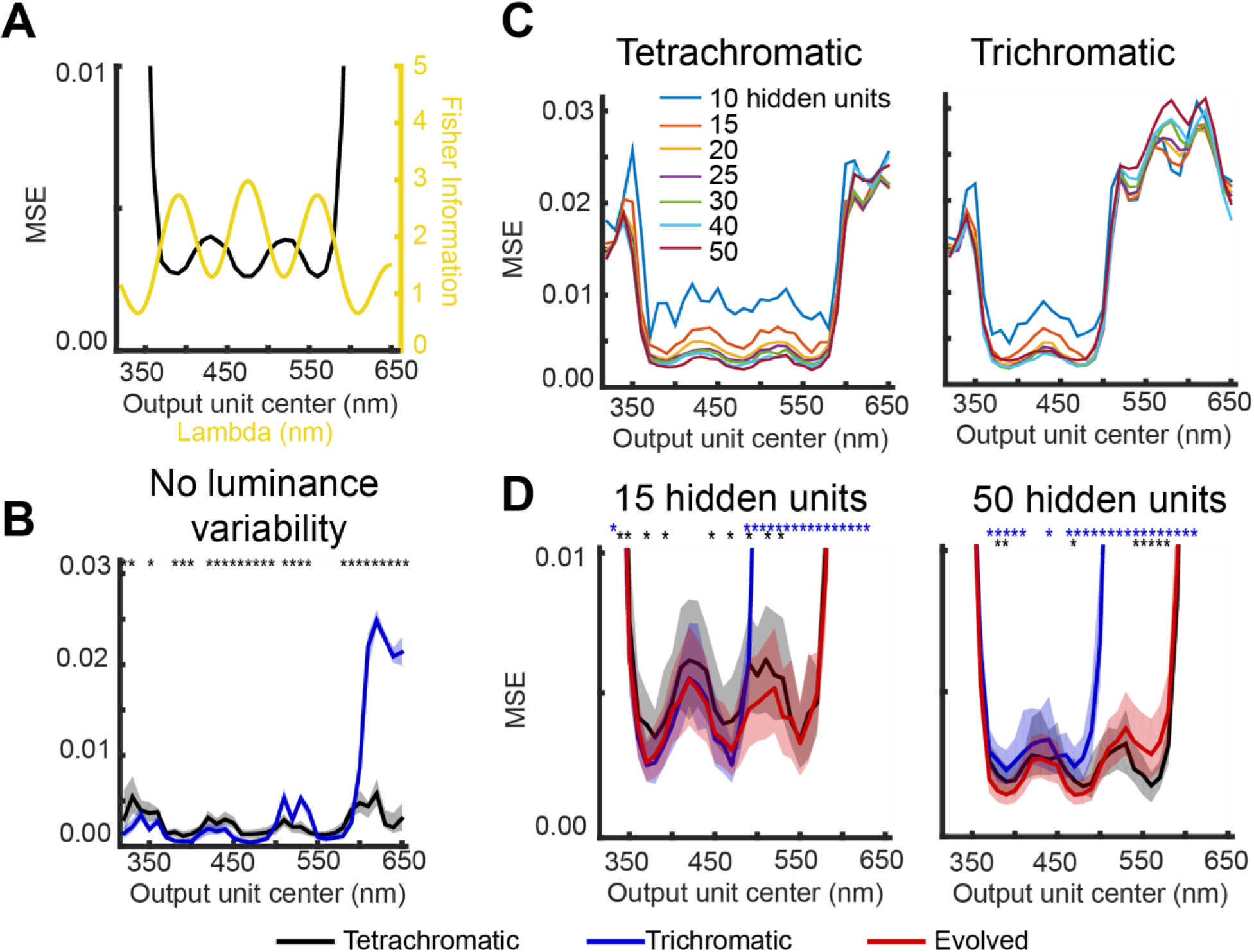
A. Black shows the performance of de novo tetrachromatic networks, showing a zoomed in version of the same plot in Fig. 1D. Yellow shows the Fisher Information of the input layer for networks with UV, blue, green, and red photoreceptors. B. Tri- and tetrachromatic networks were trained with a training set of monochromatic stimuli that lacked luminance variability. Although differences between tri- and tetrachromatic networks were observed (asterisks indicate significance with Cohen’s d > 0.5), networks perform similarly and lack clear differences for long most long wavelength output units. C. Performance of de novo networks with variable hidden layer size. Performance begins to asymptote around 15 hidden units, and regardless of hidden layer size, long wavelength output units perform poorly for trichromatic networks. D. Networks with 15 or 50 hidden units were created using the two-stage training procedure and were compared to de novo networks. Data shows the performance after 100 training epochs. Shading shows the 25^th^ and 75^th^ percentile of performance. Asterisks mark output units that were significantly different between evolved and either tri- (blue) or tetrachromatic (black) networks with Cohen’s d > 0.5.

**Figure S2.**
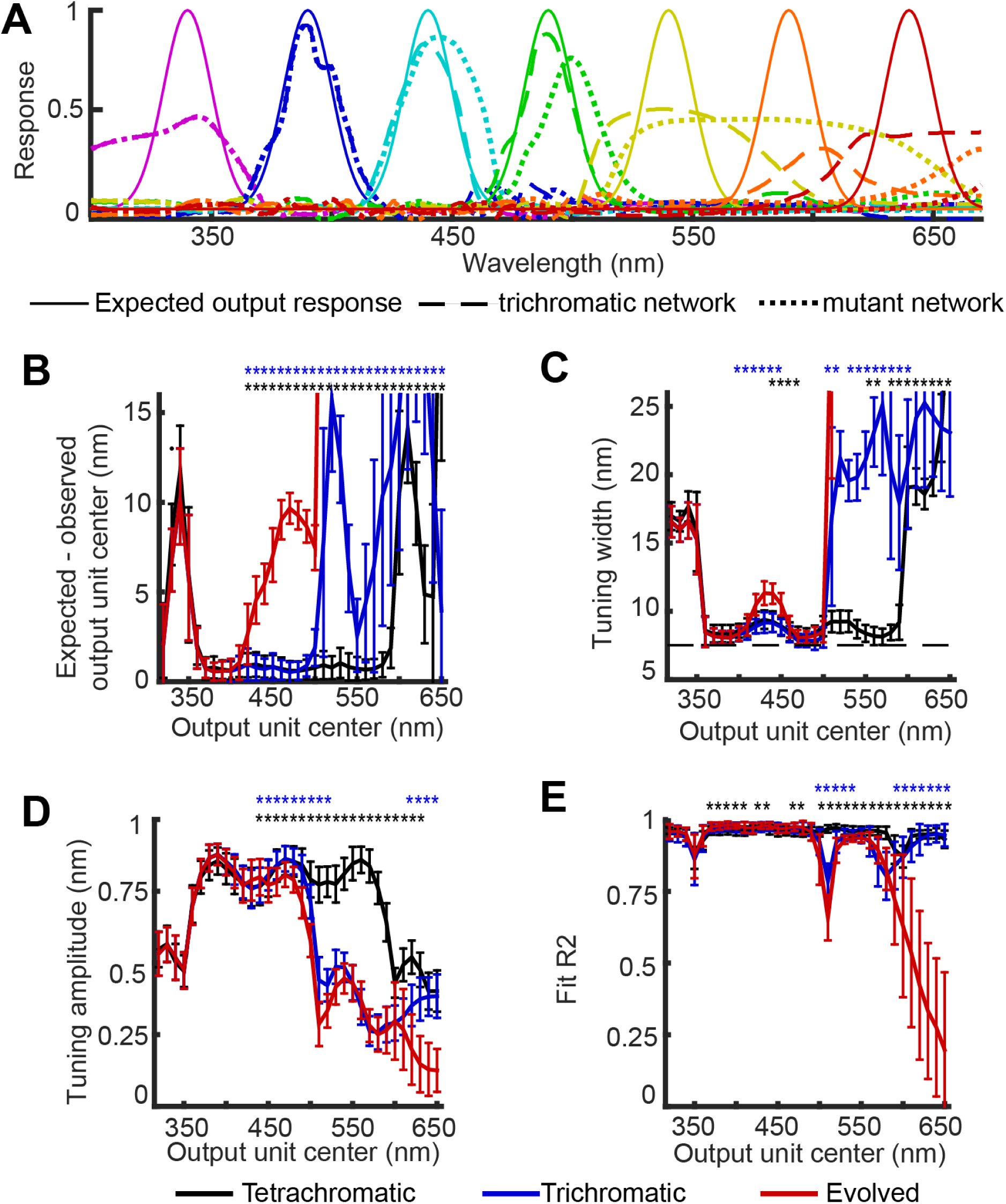
A. Example output tuning curves for a trichromatic network and the same network after mutating the input layer to tetrachromatic. B-E. Instead of calculating the MSE, we instead fit Gaussians to each output unit tuning curve. Each panel shows a comparison of the fit parameters, with blue and black asterisks indicating significant differences between evolved and de novo tri- or tetrachromatic networks, respectively, with Cohen’s d > 0.5. In panel B, the trichromatic dip at 550 nm is observed because all long wavelength output units exhibited broad tuning centered on the inflection point of the green photoreceptor. In panel C, the dotted line denotes the expected tuning width.

**Figure S3:**
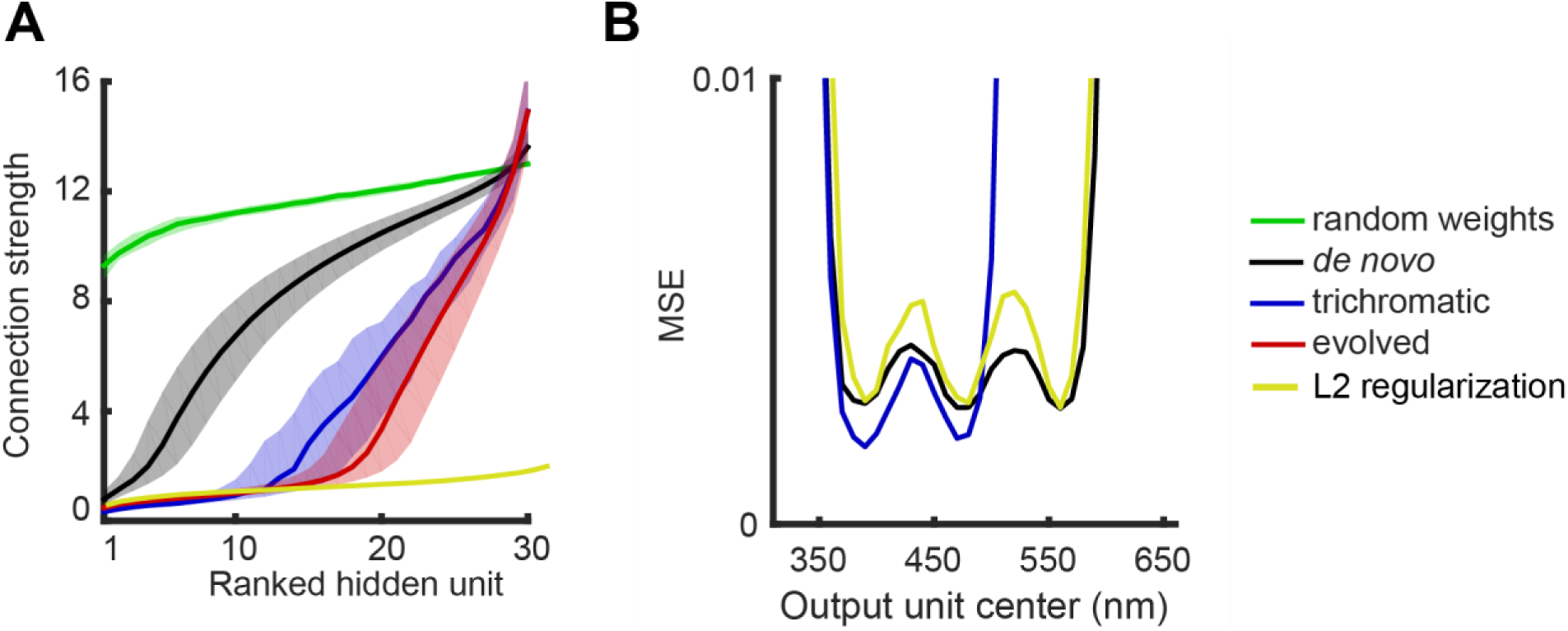
Input layer affects distribution of connection strengths. A. Connection strength was measured as the L1 norm of the input weights to a hidden unit. For each network, hidden units were ranked in order of increasing connection strength. For the yellow line, *de novo* tetrachromatic networks were trained in the same way as the black line, but the training protocol included an L2 regularization factor. Shading shows the 25^th^ and 75^th^ percentile B. L2 regularization was unable to improve the performance of *de novo* tetrachromatic networks.

**Figure S4.**
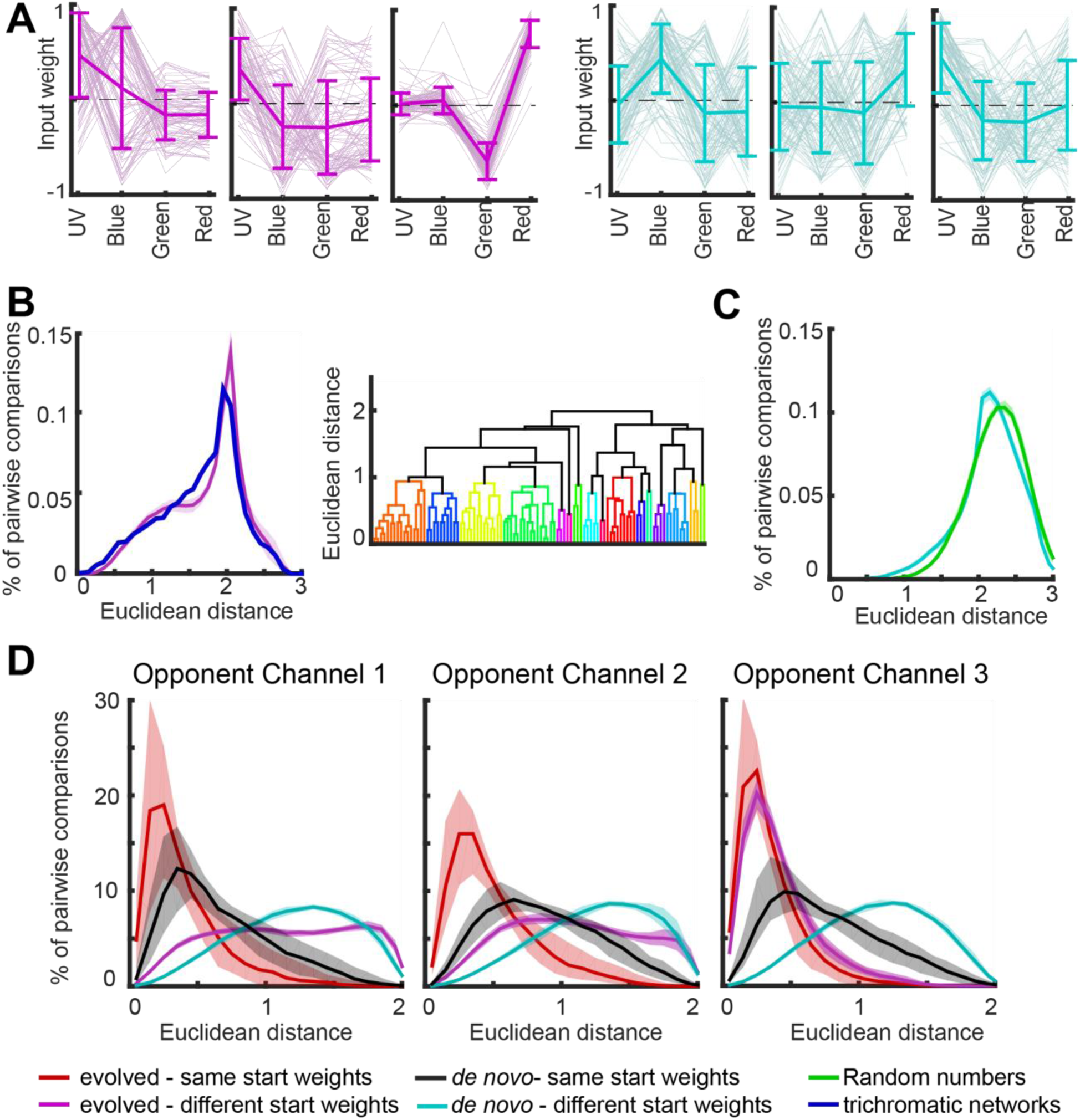
A. Example opponent channels for a group of 100 evolved (left) and de novo (right) networks that each had a different set of starting weights. Note the green vs. red opponency for channel 3 for all evolved networks regardless of the ancestral network. B. (left) The distribution of pairwise distances was computed for the trichromatic networks that were subsequently evolved and compared to groups of evolved networks with different trichromatic starting weights. (right) Dendrogram shows the computational similarities between the 100 trichromatic networks that were subsequently evolved. C. A null distribution of opponent channel similarities was created generating opponent channels using random numbers and scaling them to a unit vector to match the PCA analysis. The pairwise distance distribution is shown in comparison to the distribution for groups of de novo networks with different random starting weights. D. Opponent channel similarities were examined for each individual opponent channel rather than as the concatenated group of 3. For opponent channel 3, note the shift towards 0 for groups of evolved networks with different starting weights to overlap substantially with groups of evolved networks with the same starting weights.

**Figure S5.**
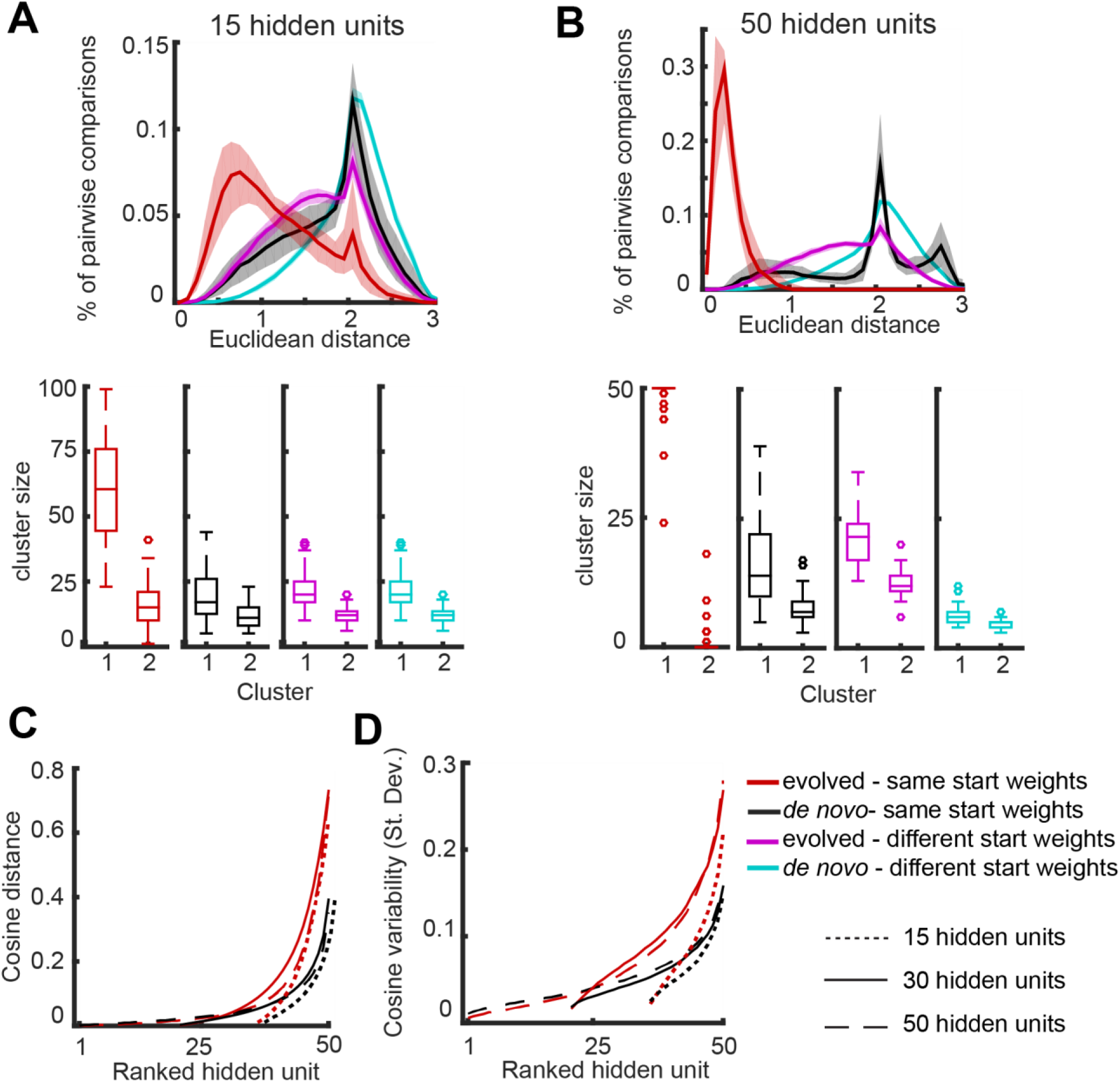
A. Opponent channels were analyzed for networks with only 15 hidden units. B. Opponent channels were analyzed for networks with 50 hidden units. Note that due to the time required to train these networks, only 50 trichromatic networks were generated and each was evolved 50 independent times. C. Cosine distances for ranked hidden units are shown for networks 15, 30, and 50 hidden units. Data are aligned so that rank 50 is the largest distance for each network, such that networks with 15 and 30 hidden units only extend to ranks 36 and 21, respectively. D. Same as C, except showing the variability in cosine distances.

**Figure S6.**
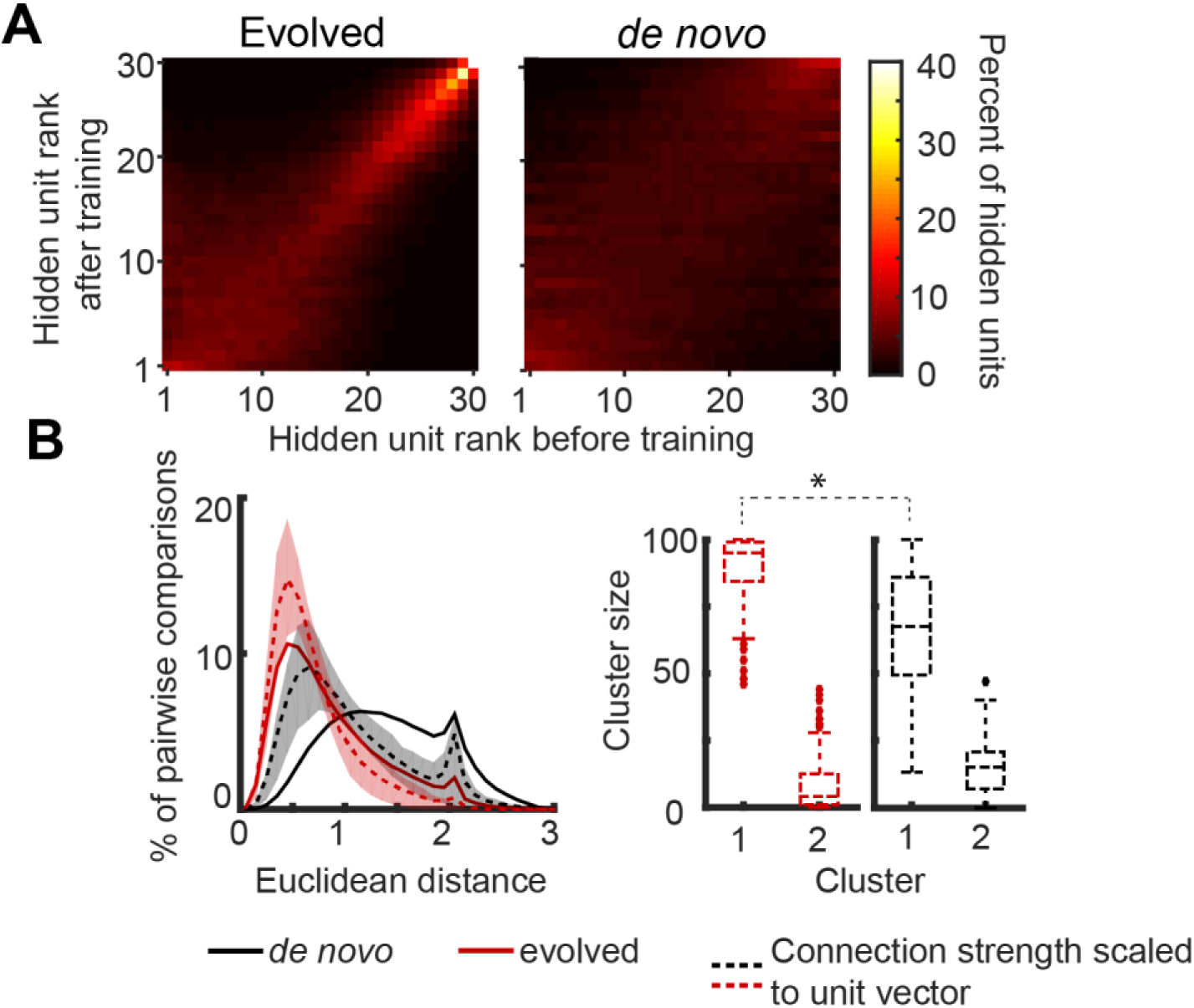
A. Connection strengths were measured as the sum of the absolute value of the input weights. This connection strength was measured for each hidden unit both before and after training. The 30 hidden units for each network were then ranked in order of increasing connection strength. Plots show how the rank of a hidden unit changed after training. Correlations between trained and untrained weights were significant for both evolved (r = 0.72 ± 0.10) and *de novo* (r = 0.43 ± 0.12) networks, with evolved significantly more correlated than *de novo* (Cohen’s d = 1.9). B. We assessed the relative contribution of opponent tuning and connection strength to the observed opponent channels by removing the influence of connection strength. The input weights of each hidden unit were scaled to a unit vector, which retains the precise opponent interactions but gives every hidden unit a connection strength of 1.0. We then reperformed the PCA analysis. The left shows the distribution of pairwise distances, and the right shows the cluster size for the two largest clusters.

